# A touch that lingers: synchronized patting strengthens neural tracking of auditory rhythm in newborns

**DOI:** 10.64898/2026.10.09.758111

**Authors:** Monica Hegde, Mohammadreza Edalati, Anne Kösem, Arthur Foulon, Stéphanie Lefebvre, Fabrice Wallois, Laurel Trainor, Barbara Tillmann, Sahar Moghimi

## Abstract

Rhythmic auditory stimulation in early life rarely occurs in isolation. Caregivers naturally combine singing and speech with rhythmic touch, rocking, and visual cues. Newborns are sensitive to auditory rhythm from birth, yet little is known about how temporally aligned information across sensory modalities shapes the neural encoding of auditory rhythmic structure. Here, we tested whether rhythmic touch can alter how the newborn brain tracks an ongoing auditory signal beyond the period of tactile stimulation. We recorded high-density electroencephalography from full-term newborns (N = 40), 1-3 days after birth, while they were exposed to recorded French lullabies presented either alone or accompanied by gentle patting synchronized with the metrically strong beat. Patting was delivered intermittently, allowing us to compare neural responses to lullabies across conditions, notably during periods without concurrent tactile stimulation. We quantified neural tracking of the lullabies using brain-stimulus coherence and temporal response function modeling. Newborns’ neural activity synchronized with the rhythmic structure of the lullabies at frequencies corresponding to both the beat and the metrically accented strong beat. Critically, neural tracking was stronger when the lullabies had been accompanied by synchronized touch than when they were presented alone, and this enhancement persisted during subsequent periods in which no touch was delivered. Although rhythmic touch enhanced tracking in newborns with both weaker and stronger auditory-only tracking, there was a larger gain in those who initially showed weaker auditory tracking. Together, our findings reveal that the newborn brain exploits rhythmic touch to strengthen its encoding of auditory rhythm, and that this benefit outlasts the touch itself. This provides neural evidence that patting an infant in time with a song, a caregiving practice shared across cultures, does more than simply provide a tactile stimulation. It shapes how the newborn brain encodes the song’s rhythm.

## Introduction

One of the brain’s fundamental capacities is to extract structure from the continuous flow of sensory information as it unfolds rapidly over time. Cross-modal interactions are crucial in this regard as they can enhance the processing of a percept that would be less accurate if based on a single sense (Atilgan et al., 2018; Morillon et al., 2014). Compelling illustrations come from speech and music, where adding sensory or motor information, such as visual cues (Alsius et al., 2016), tactile input (Guilleminot & Reichenbach, 2022; Huang et al., 2012; Răutu et al., 2026), or tapping along (Morillon et al., 2014; Morillon & Baillet, 2017), can help listeners extract the underlying rhythmic structure. Far from trivial, this capacity may be especially consequential early in life, when communication is inherently multisensory and temporally structured with facial movement, gesture, touch, and body movement accompanying speech and singing, providing infants with redundant rhythmic information across sensory modalities (Alviar et al., 2026; Bahrick et al., 2004; Streri & de Hevia, 2023). Importantly, this multisensory rhythmic stimulation is present even prenatally, as the fetus is continuously exposed to recurrent maternal rhythms, including physiological cycles (e.g., heartbeat, breathing), locomotion, and vocalizations (Lecanuet & Schaal, 1996; Provasi et al., 2014; Russo et al., 2024), which integrate fetal vestibular and somatosensory stimulation with movement-related, acoustic and vibrotactile signals. Together, these observations raise an important question regarding whether the biological predisposition in humans to exploit and integrate multisensory temporal regularities is present from the earliest stages of sensory experience to make sense of their environment.

To test this directly, we developed an experimental model in which human newborns were studied during the first three days of life using electroencephalography (EEG) to probe the human brain’s early integration of multisensory rhythmic information and its potential priming power. Auditory neural responses to sound have already been demonstrated in fetuses (Moser et al., 2021) and in premature newborns (Mahmoudzadeh et al., 2013). Regarding auditory rhythm perception in particular, EEG evidence shows that the neonatal brain forms temporal expectations while listening to music (Bianco et al., 2026), tracks auditory rhythmic hierarchies and meter-related periodicities (Edalati et al., 2023, 2024; Saadatmehr et al., 2025), and extracts the underlying beat (Winkler et al., 2009). Neonatal neural processing of auditory temporal regularities extends beyond the auditory cortex into somatosensory and premotor cortices (Mashhadi et al., 2025). Furthermore, neonatal neural evoked responses to auditory-tactile stimuli exceed the sum of unimodal responses, with this superadditivity reflecting early multisensory integration (Ronga et al., 2021). Despite this evidence, it remains unknown whether multisensory rhythmic cues can sharpen auditory rhythm perception in newborns during continuous, unfolding stimulation.

Exposure to rhythmic structures can shape adults’ perceptual processing beyond the stimulation itself, influencing the encoding of subsequently presented linguistic material (Fiveash et al., 2021; Kösem et al., 2018; Zoefel et al., 2018; Zoefel & VanRullen, 2016). EEG evidence suggests similar mechanisms in young infants, with rhythmic primes biasing the metrical perception of an ambiguous rhythm (Flaten et al., 2022). Evidence also shows persistent effects across modalities in infants, with vestibular stimulation through bouncing altering the subsequent metrical interpretation of an auditory rhythm (Phillips-Silver & Trainor, 2005, 2007). A fundamental question, however, is whether such capacities are already present at birth. Specifically, it remains unknown whether multimodal rhythmic information experienced by newborns can persist beyond the period of stimulation to influence auditory processing. Such an effect, observed before extensive postnatal experience, would suggest that the capacity of rhythmic context to organize subsequent perception is a fundamental property of the human brain.

Across cultures, caregivers instinctively pair their singing with rhythmic touch, patting, rocking, or walking (Alviar et al., 2026). Does this everyday correlated multisensory rhythmic stimulation enhance how the newborn brain tracks song? To find out, we recorded EEG from newborns while they listened to recorded lullabies with a beat at ∼2 Hz and a metrically strong beat at ∼1 Hz, presented either alone (auditory-only) or accompanied by gentle patting synchronized with the strong beat (auditory-tactile), mimicking the rhythmic touch caregivers spontaneously provide while soothing an infant. Crucially, the patting was intermittent, with tactile stimulation delivered over four strong beats, then withheld over the next four, and so on. This allowed us to quantify neural tracking of the lullaby during listening windows that were themselves free of concurrent touch. This design allowed us to test whether temporally aligned touch can have a lasting influence, enhancing the neural tracking of auditory rhythm at birth, beyond merely adding a transient somatosensory response. Because newborns have had virtually no postnatal experience with caregiver interactions in which such multisensory rhythmic cues are paired, such an effect would suggest that the capacity to integrate rhythm across the auditory and tactile senses is present early in development, rather than emerging only later following neural maturation and experience with multisensory caregiver interactions.

## Results

Caregivers across cultures combine singing with rhythmic touch when they rock, carry, or soothe their infants. Yet, it is unknown whether touch delivered in time with a song’s beat subsequently reshapes how the infant’s brain tracks the song, or whether it merely adds a somatosensory response that ends when the touch ends. To answer this question, we recorded high-density EEG from sleeping full-term newborns, 1-3 days after birth (at the onset of exposure to such multimodal rhythmic stimulation), while they listened to five recorded French lullabies, sung by a music therapist. Each lullaby carried a beat at ∼2 Hz and a metrically accented strong beat at ∼1 Hz, and was presented in two conditions: auditory-only (AO), in which newborns simply heard the lullabies, and auditory-tactile (AT), in which the experimenter gently patted the newborn’s left shoulder in synchrony with the 1 Hz strong beat (Fig. 1A). Patting was realized in repeating 8-s cycles, corresponding to eight strong beats of the lullaby: patting systematically accompanied the first four strong beats (4 s) and were withheld during the next four (4 s; Fig. 1B). Patting was thus delivered only during the second half of each 8-s cycle, so that neural responses to the lullaby could be compared between conditions during windows free of concurrent tactile input. We evaluated how the neural activity tracked the lullabies’ acoustic temporal dynamics, quantified as either spectral flux or envelope, at two levels of analysis: brain-stimulus coherence and temporal response function (TRF) modeling (Fig. 1D-E).

**Figure 1.**
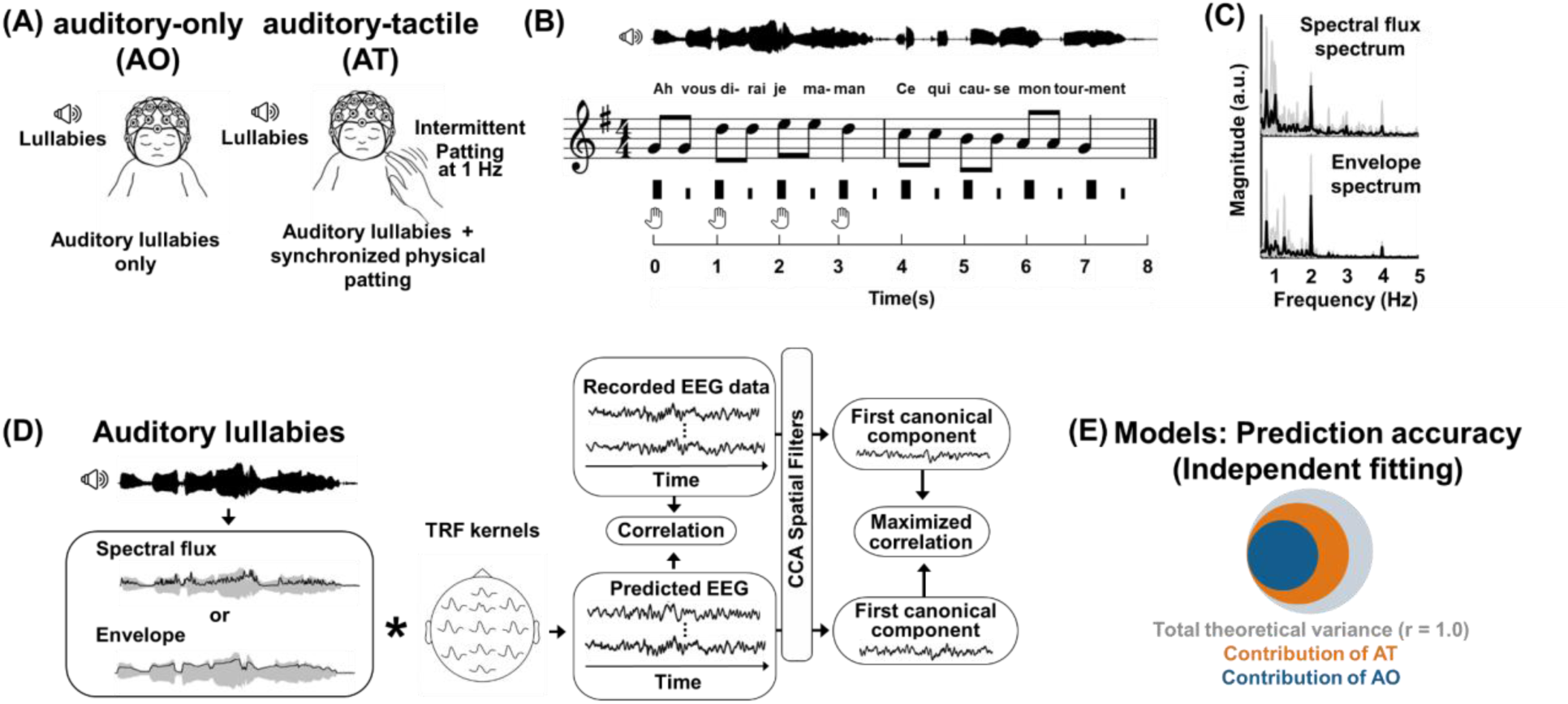
Experimental paradigm, stimulus structure, and analysis pipeline. **(A)** Newborns listened to recorded lullabies in two conditions: an auditory-only condition (AO) and an auditory-tactile condition (AT), in which gentle patting was delivered at 1 Hz in synchrony with the metrically strong beat of the lullaby. **(B)** Example 8-s stimulus cycle showing the acoustic waveform, musical score, beat structure, and timing of tactile stimulation in the AT condition. Patting was delivered during four consecutive strong beats and withheld during the following four beats, allowing neural responses to be assessed during periods without concurrent tactile input. **(C)** Spectral flux and envelope spectra across the lullabies, showing prominent rhythmic structure around 1 and 2 Hz, corresponding to the metrically strong beat and beat frequencies, respectively. Individual lullabies are shown in gray and the across-lullaby average in black. **(D)** Schematic of the temporal response function (TRF) and canonical correlation analysis (CCA) pipeline. Spectral flux or the envelope was used as the acoustic predictor to estimate TRF kernels and generate predicted EEG responses. Prediction accuracy was quantified by comparing predicted and recorded EEG. CCA spatial filters were subsequently applied to the predicted and recorded multichannel EEG to obtain their maximized canonical correlation. **(E)** Schematic illustration of prediction accuracy for independently fitted AO and AT models, with the contribution of each condition represented relative to the total theoretical correlation (r = 1).

### Newborn’s brain tracks the auditory rhythmic structure of the lullabies

To investigate whether neural activity followed the rhythmic structure of the lullabies (Fig. 1C), magnitude-squared coherence was computed between the EEG data at each electrode and the spectral flux or the envelope of the corresponding lullaby, over the 8-s cycles of the AO condition across the 0.5–5 Hz frequency range. Cluster-based permutation analysis against surrogate data identified significant clusters extending over neighboring electrodes and adjacent frequency bins. More precisely, brain-stimulus coherence with spectral flux showed two significant clusters (Fig. 2A): one spanning 0.75–1.25 Hz (z-sum = 337.043, p < .001) and another one spanning 1.75–2.25 Hz (z-sum = 320.171, p < .001). The same analysis with the envelope yielded closely matching clusters (Fig. S1A): 0.75–1.25 Hz (z-sum = 375.364, p < .001) and 1.75–2.25 Hz (z-sum = 370.718, p < .001). Thus, within days of birth, the slow neural activity tracked the rhythmic structure of the sung lullabies, notably characterized by a metrical structure with the beat at ∼2 Hz and the metrically accented strong beat at ∼1 Hz. No other significant cluster emerged at other frequencies, demonstrating the specificity of the neural responses (p > .339). Finally, directly comparing coherence values obtained using the two acoustic features (spectral flux and envelope) revealed no significant clusters (p > .214).

**Figure 2.**
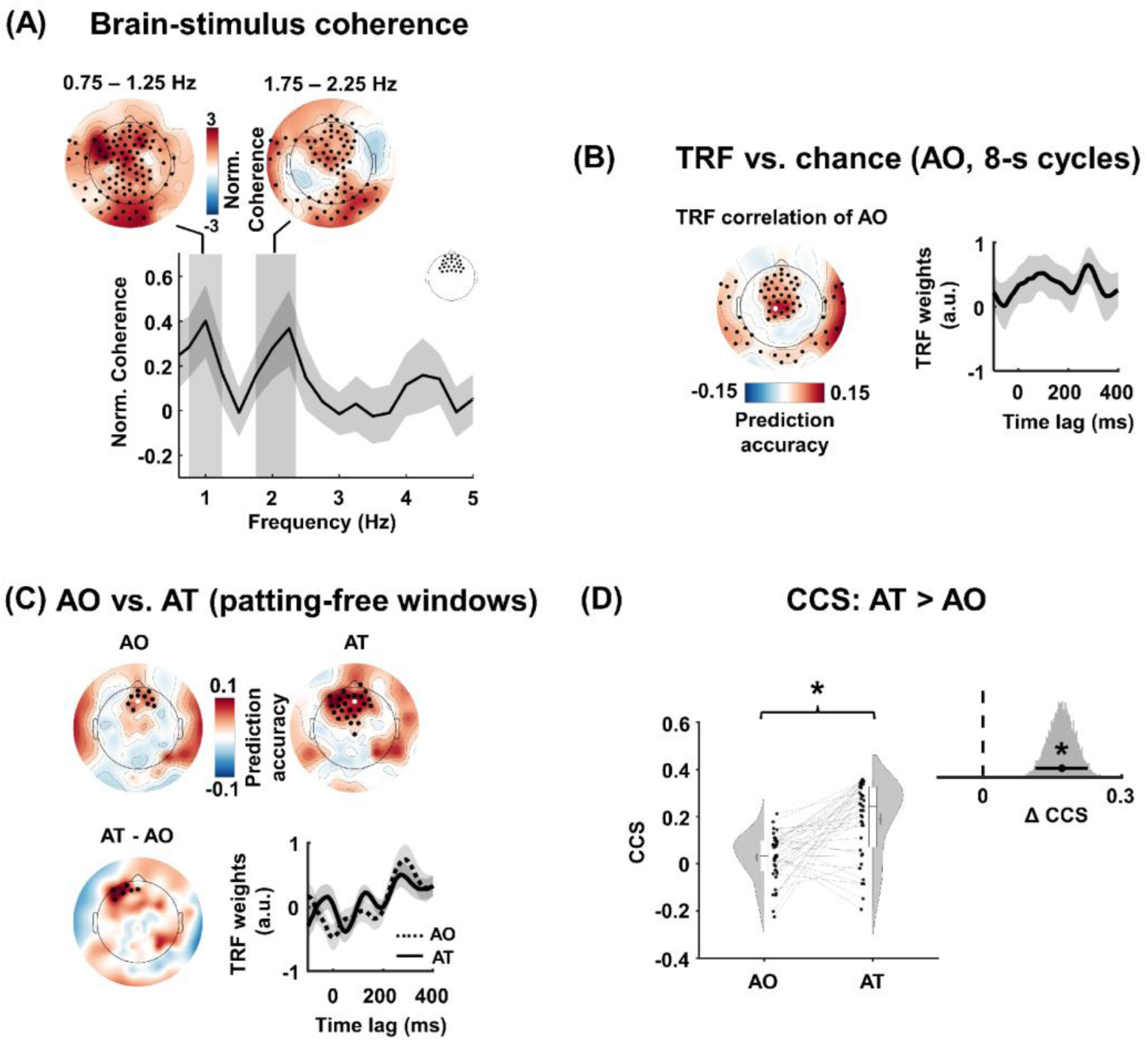
Neural tracking of the lullabies and its enhancement by tactile stimulation. **(A)** Brain-stimulus coherence between the EEG and the spectral flux in the AO condition. Topographies show surrogate-normalized coherence averaged over the two significant electrode × frequency clusters (0.75–1.25 Hz and 1.75–2.25 Hz); black dots mark the electrodes belonging to each cluster. Bottom: normalized coherence spectrum averaged over the frontal electrodes shown in the inset; solid line and shaded band show the mean and SEM across participants, and the gray vertical bands mark the frequency extent of the two significant clusters. **(B)** TRF prediction accuracy of the spectral flux model in the AO condition, computed on the 8-s cycles. The topography shows prediction accuracy, with black dots marking the electrodes of the significant cluster relative to chance; right, TRF weights (mean and SEM across participants, for the electrode marked by a white dot on the topography) as a function of time lag, peaking at about 300 ms. **(C)** Comparison of the AO and AT conditions on the tap-free windows. Top: prediction accuracy topographies per condition; black dots mark the significant clusters relative to chance. Bottom-left: AT – AO difference, with black dots marking the significant frontal cluster. Bottom-right: TRF weights over the marked electrode, with a white dot on topographies, for the AO (dashed) and AT (solid) conditions. **(D)** Canonical correlation scores per condition; gray lines connect individual newborns, half-violins show the group distributions, and the asterisk marks the significant paired difference (p < .001). Inset: distribution of the AT – AO effect posterior of β from the Bayesian regression; the black dot and horizontal bar show the mean and 94% HDI, the dashed line marks zero, and the asterisk marks p < .001.

Next, we quantified neural tracking of the lullabies’ continuous acoustic dynamics, and asked which acoustic feature the newborn EEG followed more closely. To this end, separate forward TRF models were trained with spectral flux and envelope as input, and the multichannel EEG as output. The models mapped the spectral flux or envelope, separately, onto the recorded neural response, and determined its accuracy in predicting the EEG, indexed by how strongly the brain tracked the stimulus. Models were estimated on each 8-s cycle of the AO condition. First, we compared each model’s prediction accuracy against chance level estimated from surrogate data. Cluster-based permutation correction showed two significant clusters for the spectral flux model (Fig. 2B): a frontal and frontocentral cluster (z-sum = 188.956, p < .001) and a posterior-lateral cluster (z-sum = 139.088, p < .001). The corresponding TRF weights showed that the spectral flux modulated the EEG data notably at two lags of about 100 ms and 300 ms (Fig. 2B, right), in agreement with auditory event related potentials in newborns (Edalati et al., 2022; Panzani et al., 2023). For the envelope model, the same comparison against chance showed only a frontocentral cluster (z-sum = 42.999, p = .004; Fig. S1B), with TRF weights showing a peak only at about 100 ms. Thus, both models performed better than chance. We then compared the two models directly, finding that prediction accuracy was higher for the spectral flux-based model than for the envelope-based model within a frontal and frontocentral cluster (t-sum = 95.745, p < .001; Fig. S2A). Based on this observation, we used spectral flux as the input to the TRF model for the comparison between the AO and AT conditions below. However, the corresponding results for the envelope model are also reported in the supplementary material (Fig. S2B).

### Tactile stimulation enhances neural tracking beyond the stimulation window

To investigate the effect of tactile stimulation on neural tracking after the stimulation period, we compared TRF models between the AO and AT conditions on the first 3.5 s of the patting-free half of each 8-s cycle (Fig. 1B). The final 0.5 s of the patting-free window was discarded to avoid contamination by the upcoming patting of the next cycle; the tactile sensor recordings confirmed that this window was free of patting for all retained trials (Fig. S3). Coherence was not suited for this comparison because a magnitude-squared coherence estimate requires averaging over several spectral segments, and dividing a 3.5-s window into segments short enough to average would leave a frequency resolution too coarse to isolate the 1 and 2 Hz target frequencies, leaving coherence values dominated by short-sample bias. We used TRF, which is fitted in the time domain across all windows, and therefore is not subject to this limitation. Models were estimated separately for the AO and AT conditions on identically segmented data (acquired when listening to the identical musical segment), so that the two models differed only in whether the segments had been preceded by 4 s of patting.

In both conditions (AO and AT), prediction accuracy was higher than chance over frontal electrodes (AO condition: z-sum = 35.481, p = .01; AT condition: z-sum = 86.169, p < .001; Fig. 2C, top). Comparing the prediction accuracy of the two models showed that accuracy was higher in AT than in AO within a significant frontal cluster of electrodes (t-sum = 16.321, p = .039; Fig. 2C, bottom-left). TRF weights for spectral flux to predict EEG signals in AO and AT conditions are shown in Fig. 2C. The TRF weights were similar in the two conditions, with deflections at lags of about 150 ms (more pronounced in the AT condition) and 300 ms.

We further investigated the difference between AO and AT conditions with canonical correlation analysis (CCA) applied to the TRF-predicted and recorded EEG. CCA finds a pair of spatial filters that maximizes the correlation between the two multichannel signals, pooling stimulus-driven activity distributed across electrodes into a single score per newborn and condition. This multivariate analysis confirmed the electrode-level observation: canonical correlation scores (CCS) were higher in the AT than in the AO condition (t(38) = 5.204, p < .001, d = 1.197, BF_10_ = 2800; β = 0.171, 94% HDI = [0.114, 0.227]; Fig. 2D). Together, the multivariate and univariate measures indicate that lullabies presented with intermittent synchronized touch were tracked more strongly than the same lullabies presented without touch, during listening periods that were themselves free of tactile input This suggests that the impact of gentle strong-beat patting persisted beyond the period of tactile stimulation and that tactile input enhanced the neural encoding of the lullaby rather than merely adding a concurrent somatosensory response.

### Tactile stimulation compensates for weak auditory tracking

We then asked whether this beneficial effect of tactile stimulation on rhythm tracking was uniform across the population, or whether some newborns benefitted more than others. To investigate this, we asked whether the tactile enhancement depended on a newborn’s tracking level in the AO condition. Participants were ranked based on their CCS in the AO condition and split into three groups: the 15 participants with the lowest scores formed the Low AO group, the 15 with the highest formed the High AO group, and the 9 intermediate participants were left unassigned to increase the inter-group distance (Fig. 3A). Neural tracking was stronger in the AT than AO condition within both groups: the Low AO group (t(14) = 5.07, p < .001, d = 2.031, BF_10_ = 179.219; β = 0.288, 94% HDI = [0.198, 0.382]; Fig. 3B, bottom) and High AO group (t(14) = 3.383, p = .005, d = 1.095, BF_10_ = 9.935; β = 0.12, 94% HDI = [0.06, 0.18]; Fig. 3B, top).

**Figure 3.**
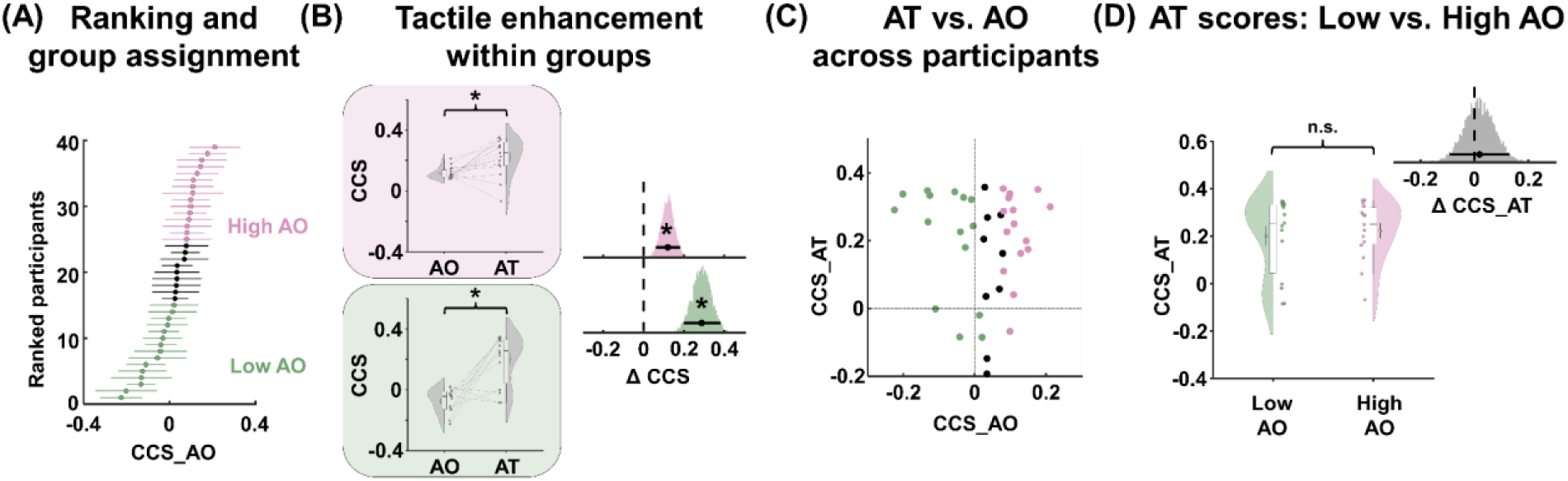
Tactile stimulation raises neural tracking to a comparable level in newborns with low and high auditory-only tracking. **(A)** Participants ranked by their canonical correlation score in the AO condition. Dots show participant means and horizontal lines show the standard error across trials; green and pink mark the Low AO and High AO groups, respectively (n = 15 each), and black marks the 9 unassigned intermediate participants. **(B)** Canonical correlation scores in the AO and AT conditions within the High AO (top, pink) and Low AO (bottom, green) groups; lines connect individual newborns and asterisks mark significant paired differences (p < .001 for Low AO and p = .005 for High AO). Right: distributions of the AT – AO effect per group posterior of β, with dots and bars showing the mean and 94% HDI and the dashed line marking zero. **(C)** AT versus AO canonical correlation scores per participant, colored by group. **(D)** Canonical correlation scores in the AT condition for the Low AO and High AO groups (difference n.s., p = .685). Inset: distribution of the between-group difference with its mean and 94% HDI; the dashed line marks zero.

Fig. 3C shows the relation between CCSs in the two conditions across participants and groups. Despite the groups’ markedly different AO scores, neural tracking of the lullabies in the AT condition did not differ between groups (t(28) = 0.41, p = .685, d = 0.15, BF_10_ = 0.367; β = 0.018, 94% HDI = [-0.092, 0.13]; Fig. 3D). Because the absence of a significant difference does not necessarily demonstrate similarity, we tested for equivalence directly with the interval Bayes factor. This tests whether the group difference falls within a margin of negligible size, which must be specified in advance. We computed the interval Bayes factor at two margin values selected based on two conventional effect-size benchmarks: a small effect with m = 0.2, and a medium effect with m = 0.5; Cohen, 2013). For CCS in the AT condition, the data from the Low and High-AO groups were ∼4 times more likely under equivalence than under a genuine group difference for a small effect (m = 0.2, BF_10_ = 3.991; moderate evidence) and ∼10 times more likely under equivalence for a medium effect (m = 0.5, BF_10_ = 10.048; strong evidence). Together, these results suggest that tactile stimulation enhanced neural tracking in both groups. However, the gain in neural tracking due to tactile stimulation was higher in the Low AO group, bringing them to a comparable level to newborns with high auditory tracking, with any group difference reliably smaller than a medium-sized effect.

## Discussion

During the first months of life, caregivers interact with their infants in an inherently multisensory way, including rhythmically coordinated rocking, touching, and speaking or singing. We showed that the newborn brain already exploits this multisensory rhythmic structure. Within a couple of days after birth, rhythmic tactile stimulation enhances newborns’ neural tracking of an auditory rhythm. Crucially, this enhancement outlasts the tactile stimulation itself, as rhythmic patting improved auditory processing beyond the period of concurrent touch. These results reveal two neural capacities present remarkably early in human ontogeny. First, cross-modal interactions sharpen the perception of temporal structure even during the very first days of life. Second, brief, ecologically natural stimulation leaves a sustained trace that continues to shape how the brain processes the auditory environment. These observations suggest that tactile rhythmic input can modulate an internal representation of a song’s periodic structure, likely by cueing the strong beat that organizes the rhythm, and increasing attention to the building blocks of the auditory rhythmic content (Henry & Herrmann, 2014; Large & Jones, 1999).

The benefit of tactile stimulation was larger in newborns whose auditory tracking was initially weakest, suggesting that redundant temporal information across the senses was most useful when auditory rhythm representation was fragile. A conceptually similar pattern is reported in children with developmental language disorder (Fiveash et al., 2023). However, a parsimonious alternative explanation is a ceiling effect (Falk et al., 2017; György et al., 2025; Ostergaard, 1998), with less room for improvement in infants whose brains strongly tracked the auditory rhythm (although the difference between conditions was still significant). Our data cannot fully separate these accounts, although the two are not mutually exclusive. In either case, it is important to note that infants with weaker tracking improved with the tactile stimulation to become indistinguishable from those with initially stronger tracking.

TRFs from continuous rhythmic stimulation may reflect overlapping stimulus-evoked responses (Nguyen et al., 2026; Prinsloo & Lalor, 2022) and neural activity related to internal representations (Bianco et al., 2024; Di Liberto et al., 2020; Di Liberto et al., 2015). In the current study, the analysis window argues against a tactile-evoked account. Thus, the near-zero-lag component may include stimulus-driven, but also importantly expectation-related activity, although the present TRF does not allow these contributions to be separated. A later response at approximately 250-300 ms falls within the latency range of sensory-evoked cortical responses in early development (Fabrizi et al., 2011; Van Der Vaart et al., 2022). Because a similar component occurred in the auditory-only condition, it cannot be considered tactile-specific. Auditory event-related potentials in neonates and young infants also contain prominent responses in this latency range (Mahmoudzadeh et al., 2016; Polver et al., 2023). The late component could therefore reflect auditory processing, overlapping auditory and somatosensory responses, or a more general response to temporal structure.

Several non-exclusive mechanisms could explain the enhancement. Rhythmic tactile stimulation on the strong beat may align slow neural activity with external temporal structure (Breska & Deouell, 2017) and increase sensitivity to temporally expected events (Fiveash et al., 2021; Large & Jones, 1999). Such effects can outlast even beyond the stimulation window (Flaten et al., 2022; Van Bree et al., 2021), consistent with neural entrainment to rhythmic information (L’Hermite & Zoefel, 2022; Zoefel & Kösem, 2024), particularly given the salient beat of the sung lullabies and the anticipatory periodicity encoding already present in preterm neonates (Edalati et al., 2024). In addition, temporally coordinated information across modalities can lead to auditory gain (Guilleminot & Reichenbach, 2022; Morillon & Baillet, 2017; Răutu et al., 2026) and increase the salience of temporal properties of early communication such as rhythm and tempo (Alviar et al., 2026; Bahrick et al., 2004). Common to these accounts is the idea that touch does not merely provide a temporal cue while it is present, but strengthens an internal representation of the lullaby’s rhythmic structure that continues to guide the tracking of the auditory stream once touch has ended. This temporal-scaffolding interpretation is also consistent with the larger benefit observed in newborns with weaker auditory-only tracking. General effects of touch on arousal or physiological state may also have contributed, and are not mutually exclusive with a rhythm-specific account. However, arousal alone seems unlikely to explain the present results, as newborns remained asleep throughout the recording, and the comparable response latencies and spatial patterns across conditions indicate that touch enhanced the same auditory tracking process rather than inducing a global change in brain state. Disentangling the respective contributions of rhythm-specific and general effects goes beyond this initial demonstration and will be an important next step, for example by comparing patting on the strong beat with regular patting on an unrelated tempo, and by monitoring physiological state alongside neural tracking.

Previous evidence suggested the involvement of sensorimotor cortical regions in auditory beat perception in preterm newborns (Mashhadi et al., 2025). The observed top-down impact of somatosensory stimulation on neural tracking of rhythm may be evidence for a developing version of the mechanisms by which sensorimotor networks impact auditory perception (Assaneo & Poeppel, 2018; Fujioka et al., 2012; Morillon & Baillet, 2017). In the newborn brain, the dorsal fiber tract from the temporal cortex to the sensorimotor cortex is already myelinated at birth (Skeide & Friederici, 2016), and primary sensorimotor cortices mature before associative cortex (Allievi et al., 2016; Gogtay et al., 2004). These early structural networks can provide the pathways for functional auditory-sensorimotor integration and the modulatory influence of somatosensory information on perception at very early stages of neurodevelopment.

Put together, this study provides electrophysiological evidence that cross modal interaction during multisensory rhythm processing is already present at birth. The lullabies we used are all constructed on a duple meter (an alteration of strong and weak beats). This same structure characterizes bipedal locomotion, and the fetus is exposed prenatally to related multisensory rhythms, such as the regular footfalls of a walking mother, which may provide a proto-musical context before birth (Larsson et al., 2019; Qirko, 2024). It is therefore possible that newborns arrive already sensitive to duple multisensory temporal organization, having encountered it in utero. A decisive test would compare multisensory enhancement across rhythmic structures that differ in their likely prenatal familiarity, contrasting duple meter against structures the fetus is less likely to have experienced.

Beyond its theoretical implications, the present finding raises the possibility that everyday multisensory caregiving could support early auditory rhythm processing. If rhythmic touch provides windows of enhanced neural tracking of auditory rhythm, as observed here, particularly in newborns with weaker auditory-only tracking, caregiving practices such as patting in time with a song might offer a natural way to scaffold temporal processing from birth, and reduce potential impairment due to atypical rhythm processing (Ladányi et al., 2020; Lense et al., 2021; Menn et al., 2022).

## Materials & Methods

### Participants

Forty-seven full-term newborns were recruited at the maternity unit of the Amiens University Hospital (CHU Amiens, France). Seven newborns were excluded because their recordings were incomplete, leaving a final sample of 40 infants (26 female). One further participant was excluded because their CSS (see Data analysis) exceeded 3 standard deviations from the group mean, leaving 39 newborns for all analyses reported in this study. Gestational age at birth ranged from 37.429 to 41.714 weeks (mean 40.026 ± 0.958 weeks). EEG was recorded in 1-3 days after birth, during sleep. No infants had a known neurological or auditory impairment. Written informed consent was obtained from a parent or legal guardian prior to participation. The ethics committee Comité de Protection des Personnes Ouest I approved the study (IDRCB: 2019-A01534-53).

### Stimuli

The auditory stimuli consisted of five familiar French children’s lullabies lasting either ∼24 s or ∼32 s (osf.io/r657v). A certified neonatal music therapist sang all lullabies while listening to a metronome at 120 bpm (2 Hz), which were recorded at 44.1 kHz. Each lullaby was therefore structured with an underlying beat at 120 BPM (2 Hz) and a metrically accented strong beat at 60 BPM (1 Hz; Fig. 1C).

Each lullaby was presented as a continuous block of six or seven consecutive repetitions, depending on its duration, so that each block lasted approximately 3 min. A period of ∼5 s of silence separated every two blocks. Stimuli were delivered through a loudspeaker positioned 1 m from the newborns’ head lying in a crib, at an average intensity of 60 dB, measured at the newborns’ head position.

### Experimental design and procedure

Each of the five blocks (one for each lullaby) was presented twice, once in an auditory-only (AO) condition and once in an auditory-tactile (AT) condition, giving 10 blocks per newborn and a total session duration of approximately 30 minutes. Block order was pseudo-randomized, with the constraints that no more than three consecutive blocks belonged to the same condition and that the same lullaby was never presented in two consecutive blocks.

In the AO condition, newborns were simply exposed to the lullabies. During the AT condition, the experimenter added rhythmic tactile stimulation by gently patting the front of the newborn’s left shoulder with her hand on a 60 BPM (1 Hz) metronome, synchronized with the strong beat and heard through headphones and audible only to the experimenter (Fig. 1A). Patting followed a repeating 8-s cycle in which four beats with patting (4-s) were always followed by four beats without patting (4-s; Fig. 1B). The patting-free half of each cycle was built into the design so that neural responses to the lullaby could be compared between conditions during periods free of concurrent tactile input.

Patting behavior of the experimenter was recorded with a sensor fixed on the front of the newborn’s left shoulder and digitized on an auxiliary channel of the EEG amplifier at 1000 Hz, ensuring sample-accurate alignment between the tactile recording and the EEG.

### EEG Acquisition and preprocessing

EEG was recorded continuously using a 124-channel HydroCel Geodesic Sensor Net (Electrical Geodesics Inc.) at a sampling rate of 1000 Hz, with a Cz vertex electrode as reference. Electrode impedances were kept below 50 kΩ. Stimulus onsets were marked in the EEG recording as TTL triggers generated by Psychtoolbox (Kleiner et al., 2007).

Preprocessing was performed in MATLAB R2022b (MathWorks) using EEGLAB (Delorme & Makeig, 2004), Fieldtrip (Oostenveld et al., 2011), APICE pipeline (Fló et al., 2022), and custom scripts. Continuous EEG data were zero-phase band-pass filtered (0.5–20 Hz) and downsampled to 500 Hz. Recordings were visually inspected, and electrodes showing persistent artifact or poor contact were removed (mean = 28.852 ± 10.05 electrodes). Independent component analysis was used to identify and remove cardiac artifacts. Next, automatic artifact detection and correction were performed with APICE. Segments in which more than 30% of channels contained artifact were rejected. Rejected electrodes were interpolated using neighboring electrodes, and the data were re-referenced to the common average. Finally, data were additionally low-pass filtered at 10 Hz and resampled to 100 Hz. Patting accuracy was also visually inspected, and 8-s cycles containing inaccurate patting (not on the beat) were marked as bad segments and excluded from further analyses. On average, 78.045% ± 8.755% of AO condition and 67.488% ± 19.022% of AT condition remained after preprocessing.

To increase the signal-to-noise ratio, EEG data were averaged across the repetitions of each lullaby within each condition. Because lullaby durations were multiples of the 8-s cycle, patting and patting-free windows occupied identical positions in every repetition, so this averaging preserved the temporal structure of the stimulation. The averaged data comprised 144 s of EEG per condition, which were segmented into 18 eight-second segments. After exclusion of segments affected by artifacts or inaccurate patting, 17.05 ± 2.36 (AO) and 16.77 ± 2.39 (AT) segments per newborn remained for analysis.

### Stimulus analysis

The temporal structure of each lullaby was quantified using both the envelope (Gehmacher et al., 2024) and spectral novelty (spectral flux; Weineck et al., 2022). The envelope was based on gamma-tone spectrograms, which were calculated over 256 frequencies, covering a range of 20 – 5000 Hz in equivalent rectangular bandwidth space (Heeris, 2018), resampled to 100 Hz, and scaled with exponent 0.6 (Biesmans et al., 2016). The spectral flux, a measure of frame-to-frame spectral change that indexes acoustic onsets and has been widely used to characterize rhythmicity in music and speech (Giroud et al., 2024; Müller, 2015; Weineck et al., 2022), was computed by converting the audio waveform to a log-magnitude spectrogram (window of 30 ms, hop size of 10 ms). The difference across successive frames was computed within each frequency bin and summed across bins, yielding a flux signal sampled at 100 Hz. The spectral content of the envelope, flux signals and the target frequencies (1 and 2 Hz) are reported in Fig. 1C.

### Brain-auditory stimulus coherence

To verify that neural activity tracked the auditory rhythmic structure of the stimuli, magnitude-squared coherence was computed between each EEG electrode and, separately, the stimulus spectral flux and envelope, using Welch’s method (MATLAB mscohere, 4-s windows, 50% overlap, frequency resolution 0.25 Hz) on the 8-s segments of the AO condition, and assessed over the 0.5–5 Hz range. Coherence was not computed for the AT condition, because the tactile stimulation was itself rhythmic at 1 Hz and would contribute to coherence at the target frequencies. Coherence was also restricted to the 8-s segments, because the 3.5-s patting-free windows used for the TRF comparison between conditions were too short for a reliable Welch estimate at the target frequencies.

### Temporal response function

To quantify neural tracking of the lullabies’ temporal acoustic dynamics, forward TRFs were estimated with mTRF-Toolbox (Crosse et al., 2016). The auditory predictor was the spectral flux (or envelope) of the corresponding lullaby, and the response was the multichannel EEG. Both the predictor and response were normalized by their standard deviation.

Two trial definitions were used. For the comparison of acoustic features (spectral flux vs envelope) within the AO condition, trials were the 8-s cycles. For the comparison between the AO and AT conditions, trials were defined as the second 4-s window of each 8-s cycle (corresponding to the 4-s patting-free window of each 8-s cycle in the AT condition). For each 4-s trial, the first 3.5 s were retained, and the final 0.5 s was discarded to avoid contamination by the upcoming patting event (Fig. S3). Identical segmentation was applied to the AO condition so that trials were matched across conditions in duration and in position relative to the lullabies’ temporal structure.

TRF models used time-lagged versions of the spectral flux (or envelope) to predict EEG activity over a lag window from –200 to 500 ms. Ridge regression was used for regularization, with λ spanning 10^−4^ – 10^8^ in logarithmic steps. The regularization parameter λ was selected by a nested cross-validation procedure. Within each outer leave-one-trial-out fold, an inner leave-one-trial-out loop over the n-1 training trials identified the λ maximizing prediction performance, which was then applied to the held-out trial.

For each participant, TRFs were estimated separately for the AO and AT conditions using the same auditory predictors (spectral flux or envelope), allowing us to test whether tactile stimulation modulated the neural encoding of the lullaby’s acoustic dynamics. Prediction accuracy was quantified as the Pearson correlation between predicted and recorded EEG on held-out trials, computed per electrode and per trial, Fisher z-transformed, and then averaged across trials.

### Canonical correlation analysis

To further assess the correspondence between the stimulus-driven EEG response and the recorded neural signal, CCA was applied to the TRF-predicted and recorded EEG. CCA identifies pairs of linear combinations of electrodes that maximize the correlation between these two multichannel signals, and therefore captures distributed response patterns that electrode-wise prediction accuracy may miss. This combination of TRF prediction and CCA-based evaluation corresponds to the CPA (canonical prediction alignment) framework introduced by Di Liberto (2026); we refer to the resulting metric as the CCS throughout.

To avoid circularity, all quantities were estimated and evaluated on disjoint data within a leave-one-trial-out scheme. On each fold, the TRF was fitted on the training trials and applied to the left-out trial to generate the predicted EEG. The canonical weights were estimated from the predicted and recorded EEG of the training trials only and then applied unchanged to the left-out trial. CCS was defined as the correlation of the first canonical component on the held-out trials, Fisher z-transformed, and averaged across trials. CCA was regularized by adding a ridge term to the covariance matrices (λ_CCA_ spanning 10^1^ to 10^5^ in logarithmic steps, selected within the training folds of the same nested scheme) to stabilize the covariance estimates given the 124-channel dimensionality.

## Statistical analysis

Statistical analyses were performed in MATLAB and Python using custom functions. Normality was assessed with the Lilliefors test when required (osf.io/r657v).

Because coherence estimates are positively biased, raw coherence values were not tested directly. Instead, a null distribution was constructed by circularly shifting the spectral flux (or envelope) relative to the EEG by a random lag and recomputing coherence, 500 times per electrode. Coherence values were expressed as a z-score relative to this surrogate distribution, and significance at each electrode and frequency of interest was assessed with the complementary error function method (Theiler et al., 1992). Family-wise error was controlled with cluster-based permutation correction across the electrode × frequency space (Maris & Oostenveld, 2007). Clusters were formed on each of 5000 permutations; one surrogate dataset was drawn per participant, passed through the identical z-scoring, group-level aggregation, and clustering pipeline, and the maximum cluster statistic across the whole space was retained to form the null distribution. The initial threshold for cluster definition and the minimum number of neighbors were set to p < .05 and two, respectively. Cluster-level statistics were reported as the sum of surrogate-normalized z-scores over the electrode × frequency points in each cluster (z-sum), along with the corresponding cluster-level p-value.

TRF prediction accuracy in each condition was compared with chance level using the same logic: surrogate prediction accuracies were obtained by circularly shifting the predictor relative to the EEG and re-evaluating the models, and observed accuracies were tested against them with cluster-based permutation correction across electrodes. Cluster statistics for these tests are reported as z-sums. TRF prediction accuracy was compared between two models or conditions with a cluster-based permutation test (Maris & Oostenveld, 2007), which controls the family-wise error rate across the 124 electrodes. The initial threshold for cluster definition and the minimum number of neighbors were set to p < .05 and two, respectively.

CCSs were compared between conditions with a paired t-test and between groups with an independent-samples t-test. In addition, each comparison was re-estimated with a Bayesian regression model fitted with Bambi (Capretto et al., 2022). CCS was modeled as a function of condition (within-participant comparisons, with participant as a group-level intercept) or group (between-participant comparisons), using the package’s default weakly informative priors. We report the posterior mean of the corresponding coefficient (β) and its 94% highest density interval (HDI). Because a non-significant difference test cannot support the absence of an effect, evidence for equivalence between groups was quantified with the interval Bayes factor (Linde et al., 2023; Morey & Rouder, 2011), using default priors (zero-centered Cauchy on effect size, scale 0.707; Rouder et al., 2009). This test compares H_1_: |δ| < m against H_0_: |δ| ≥ m, where δ is the standardized group difference, and m is the equivalence margin. An equivalence test requires specifying a margin m, i.e. the largest group difference that would still be considered negligible. We tested both m = 0.2 and m = 0.5, the conventional benchmarks for a small and medium effect, respectively (Cohen, 2013). Bayes factors were interpreted following Lee & Wagenmakers (2013), with BF_10_ > 3 and > 10 taken as moderate and strong evidence, respectively.

Differences in between-participant variances across the paired conditions were tested with the Pitman-Morgan test for correlated variances (Morgan, 1939; Pitman, 1939). Effect sizes are reported as Cohen’s d alongside Bayes factors (BF_10_) computed using the BayesFactor toolbox in MATLAB (https://github.com/klabhub/bayesFactor).

**Figure S1.**
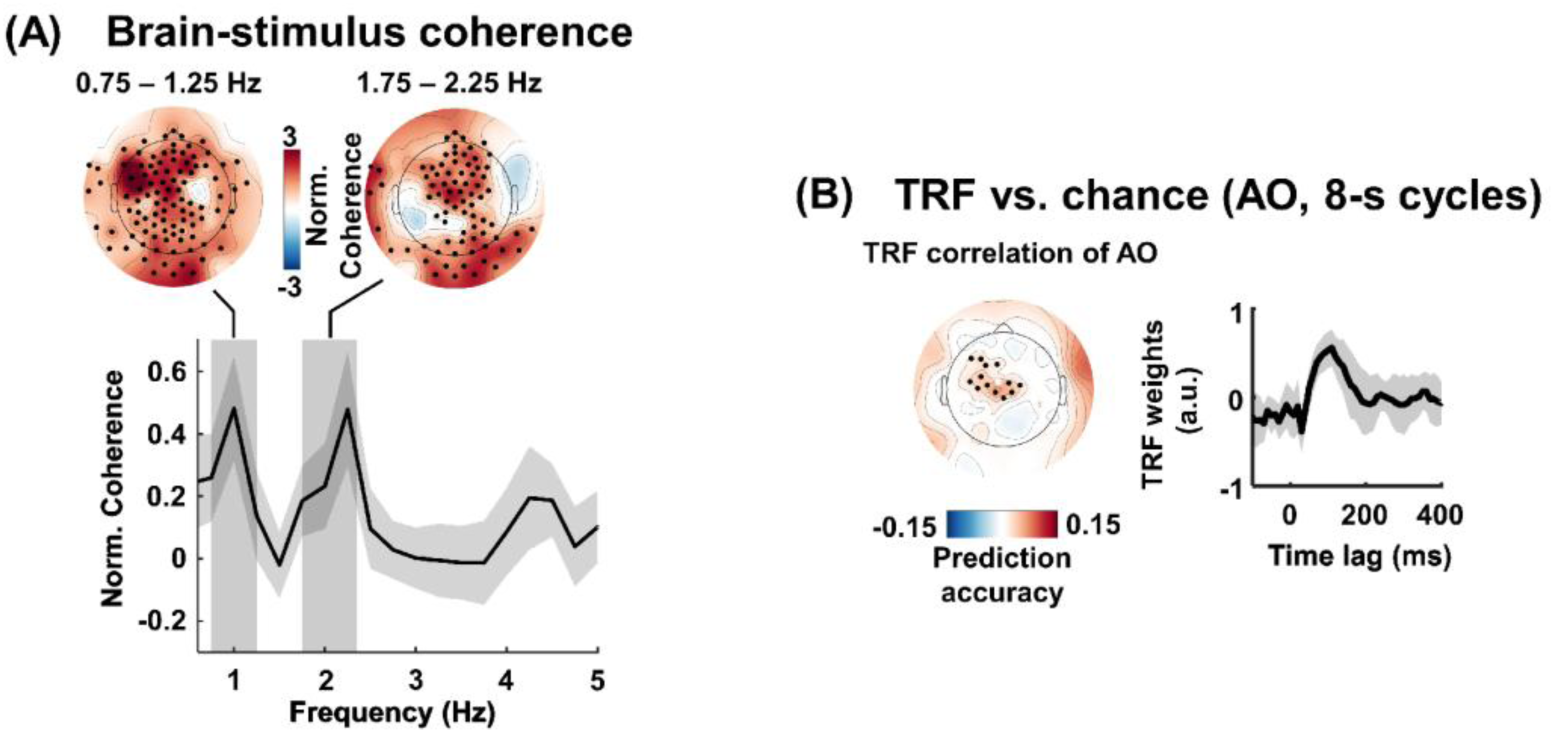
Neural tracking of the lullabies’ envelope. **(A)** Brain-stimulus coherence between the EEG and the envelope in the AO condition, as in Fig. 2A: topographies of the two significant clusters (0.75–1.25 Hz and 1.75–2.25 Hz, black dots marking cluster electrodes) and the normalized coherence spectrum with the cluster frequency ranges shaded. **(B)** TRF prediction accuracy of the envelope model in the AO condition on the 8-s cycles; black dots mark the significant frontocentral cluster relative to chance. Right: TRF weights over the cluster electrodes, peaking at about 100 ms.

**Figure S2.**
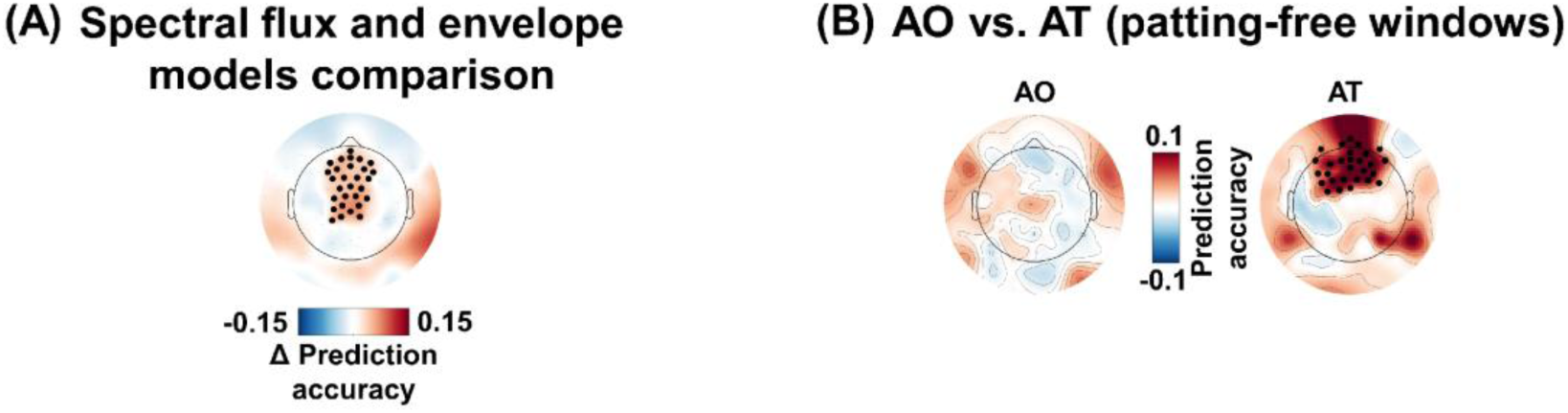
Comparison of the different models. **(A)** Topography of the difference in TRF prediction accuracy (spectral flux – envelope) on the 8-s cycles; black dots mark the significant frontal and frontocentral cluster in which the spectral flux model predicted the EEG better than the envelope model (p < .001). **(B)** Prediction accuracy of the envelope model in the AO and AT conditions on the tap-free windows; black dots mark the significant cluster relative to chance in the AT condition. There is no significant cluster relative to chance in the AO condition.

**Figure S3.**
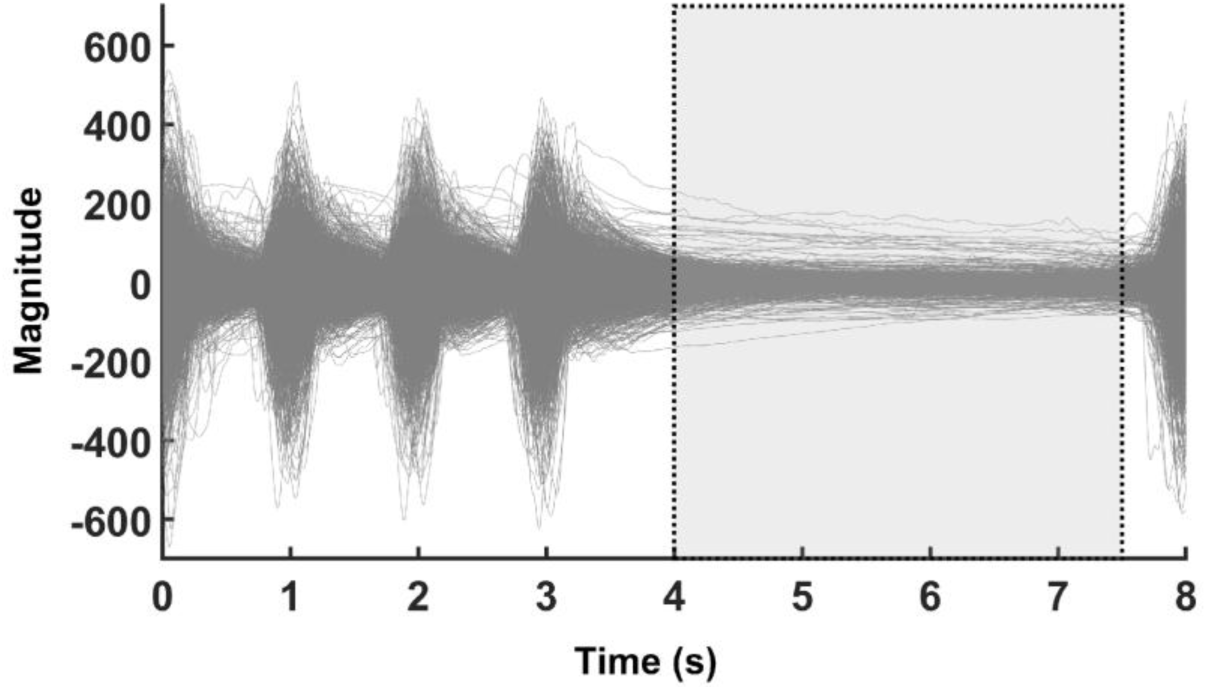
Tactile sensor recordings across the 8-s stimulation cycle in the AT condition. Gray traces show the sensor signal for all recorded cycles of all participants, aligned to cycle onset. The four taps occupy the first half of the cycle (0 – 4 s), and the dotted rectangle marks the 4 – 7.5 s analysis window used for the TRF comparison between conditions, confirming the absence of tactile input during this window. The final 0.5 s of the tap-free half was discarded because of the upcoming tap of the next cycle.

